# Spatial Sampling and Temporal Dynamics Shape Motor Decoding from Broadband Intracortical Signals

**DOI:** 10.64898/2026.09.29.755257

**Authors:** Cong Zheng, Yongxiang Xiao, He Cui

**Author notes:** Corresponding authors: Cong Zheng and He Cui.

## Abstract

**Objective:** Neural decoding performance depends strongly on neural signal representation; however, how the informative components of broadband intracortical activity vary with electrode configuration and decoding strategy remains unclear. We systematically compared the neural features spanning multiple temporal scales to identify the factors underlying robust motor decoding. *Approach.* Neural recordings were obtained from the dorsal premotor cortex, primary motor cortex, and dorsolateral prefrontal cortex of rhesus monkeys performing delayed-reaching tasks toward static or moving targets. Signals were recorded using 64-channel linear S-probes or 96-channel planar Utah arrays. We compared spiking-band power (SBP), local motor potential (LMP), and local field potential (LFP) power across multiple frequency bands using linear and nonlinear decoders. We further characterized the spectral and temporal components contributing to LMP decoding and evaluated the channel redundancy. *Main results*. The decoding performance of the SBP and high-frequency LFP features scaled with the number of recorded single units, whereas the LMP performance was more strongly influenced by the electrode geometry. LMP achieved a higher decoding performance with planar Utah arrays than with linear S-probes, accompanied by lower inter-electrode correlations and greater variability in preferred directions. Spectral analysis showed that LMP decoding was dominated by low-frequency components overlapping the delta band and movement-related cortical potentials, with an optimal decoding range of approximately 0.1–8 Hz. Nonlinear temporal decoders, particularly the convolutional neural network and gated recurrent unit models, further improved LMP decoding. *Significance*.

These findings demonstrate that the neural features most informative for motor decoding depend on both the spatial sampling configuration and the temporal structure of the recorded activity. The results highlight the importance of low-frequency population dynamics and provide principles for optimizing neural-signal representations and reducing redundant neural data for scalable and wireless brain-computer interfaces.

## 1. Introduction

The performance of brain-computer interfaces (BCIs) critically depends on the choice of neural signals. Motor-related information is embedded across a broad range of temporal scales, from slow cortical potentials and local field potential (LFP) to spiking activity and high-frequency population dynamics (Schwartz, 2007; Stark and Abeles, 2007; Pesaran *et al*., 2018). Different signal representations capture distinct aspects of the underlying neural activity and can therefore vary substantially in their relationship with behavior and their utility for decoding (Gallego-Carracedo *et al*., 2022; Abbaspourazad *et al*., 2024). Given the vulnerability of spiking activity to long-term signal instability, neural features extracted from broadband intracortical signals have emerged as alternatives to threshold-crossing rates (TCRs). Spiking-band power (SBP) is a compact high-frequency neural feature typically derived by quantifying power in the 300–1000 Hz range. Nason et al. (2020) showed that SBP closely reflects local neuronal activity and supports motor decoding comparable to or better than TCRs (Nason *et al*., 2020). Consistently, high-frequency LFP band power containing spiking activity also supports high-performance motor decoding (Irwin *et al*., 2015; Gallego-Carracedo *et al*., 2022). In contrast, local motor potential (LMP) is a low-frequency neural feature derived through low-pass filtering and temporal smoothing. Schalk et al. (2007) introduced LMP as a neural signal containing movement-related information for motor decoding, and Stavisky et al. (2015) further demonstrated that LMP alone could support fast and accurate closed-loop BCI control, while combining LMP with spikes improved performance when spike quality was poor. These findings highlight LMP as a robust, low-bandwidth neural feature with the potential for long-term and low-power BCIs (Schalk *et al*., 2007; Stavisky *et al*., 2015). However, compared to SBP, whose decoding contribution is clearly linked to spiking activity, the use of LMP has been limited by its inconsistent decoding performance across studies, potentially because its underlying neural basis remains poorly understood (Kubánek *et al*., 2009; Ince *et al*., 2010; Gunduz *et al*., 2016; Flint *et al*., 2017).

Neural activity is organized across multiple spatial scales, from cortical layers and columns to broader population-level networks, with distinct patterns of neuronal activity, synaptic inputs, and local circuit interactions across these spatial domains (Graziano, Taylor and Moore, 2002; Weiler *et al*., 2008). Extracellular signals further integrate these neural processes over spatially distinct scales depending on the biophysical properties of the recorded activity and its spatial organization (Buzsáki, Anastassiou and Koch, 2012). Consequently, the neural features that provide effective behavioral information may depend on the spatial sampling of neural activity, including cortical location, recording depth, and electrode configuration (Schwartz, 2007; Jackson and Hall, 2017). This consideration is particularly relevant for intracortical recordings, where linear probes and planar arrays, whose electrodes are distributed across cortical depth or along the cortical surface, sample neuronal populations with different spatial arrangements and sampling geometries (Maynard, Nordhausen and Normann, 1997; Mendoza-Halliday *et al*., 2024).

Therefore, systematically determining how these sampling configurations influence the neural features available for behavioral decoding is important for understanding the relationship between cortical organization, neural signal representation, and decoding performance.

Here, we systematically characterized neural signals across temporal and spatial scales for motor decoding using recordings from the dorsal premotor cortex (PMd), primary motor cortex (M1), and dorsolateral prefrontal cortex (dlPFC) of rhesus monkeys (*Macaca mulatta*). Neural activity was recorded with either 64-channel linear S-probes or 96-channel planar Utah arrays during delayed-reaching tasks toward static or moving targets. We compared conventional broadband-derived features, including SBP, LMP, and LFP power across multiple frequency bands, to determine how their decoding performance depends on the electrode geometry, frequency content, and decoder architecture. We further characterized the spectral and temporal components underlying LMP decoding and examined whether channel redundancy and nonlinear temporal structure could be exploited to improve decoding efficiency. This systematic comparison provides a framework for linking the neural basis of behaviorally relevant signals to their suitability for robust and efficient neural decoding.

## 2. Methods

### 2.1. Subjects and surgery

This study involved four adult male rhesus monkeys (*Macaca mulatta*): monkey B (7.8 kg, 4 years old) and monkey Lb (9 kg, 4 years old) implanted with 96- or 64-channel Utah microelectrode arrays (Blackrock Neurotech, USA), and monkey G (8.4 kg, 13 years old) and monkey Lp (10.0 kg, 6 years old) implanted with an acrylic recording chamber for 64-channel S-probe (Plexon Inc.) acute recording. For monkeys B and Lb, following anesthesia and stereotaxic positioning, a craniotomy was performed over the target cortical region, and the dura was opened to expose the cortical surface. The array was positioned perpendicular to the cortical surface and was gently inserted into the cortex. For monkey B, a 96-channel array was placed on the primary motor cortex (M1) and the other on the dorsal premotor cortex (PMd). For monkey Lb, an additional 64-channel array was implanted in the dorsolateral prefrontal cortex (dlPFC). The array connector was secured to the skull using dental acrylic and bone screws and the surgical site was closed.

Neural recordings began after an appropriate postoperative recovery period. For monkeys G and Lp, following anesthesia and stereotaxic positioning, a craniotomy was performed over the M1 and PMd, and an acrylic recording chamber (19 mm inner diameter) was secured over the cranial window using dental acrylic and bone screws. All the targeted brain areas were contralateral to the limbs used for reaching. All procedures have been approved by the Biomedical Research Ethics Committee of Shanghai Institutes for Biological Sciences, Chinese Academy of Sciences (approval no. ER-SIBS-221603P, version 20160310) and the Institutional Animal Care and Use Committee of Chinese Institute of Brain Research, Beijing (approval no. CIBR-IACUC-059 and CIBR-IACUC-083). They comply with national and local laws and regulations in China and are in accordance with the Guide for Care and Use of Laboratory Animals of the Institute for Laboratory Animal Research.

### 2.2. Paradigm and recording

In the manual delayed interception task, the monkey was seated in a primate chair facing a vertical touch screen, with the head restrained. Each trial began with a 600-ms hold on a central target, followed by the appearance of a peripheral target that was either static or moved along a circular path at a constant clockwise (-) or counterclockwise (+) angular velocity. After a delay, the central target disappeared (go cue, GO), instructing the monkey to initiate reach (movement onset, MO) toward the peripheral target within 800 ms. Successful trials required the touch point to fall within the predefined tolerance and be held for at least 600 ms, resulting in a liquid reward. Neural signals were acquired using the Cerebus acquisition system (Blackrock Microsystems) at a sampling rate of 30 kHz with 16-bit resolution. For each recording channel, the continuous broadband signal was simultaneously recorded for offline analysis. Task control and data acquisition were performed using Psychopy or MonkeyLogic. Hand trajectories were recorded at 100 Hz using a Vicon motion capture system (Oxford, UK) with an infrared reflective marker attached to the hand. The task parameters are presented in Table 1. Data from monkeys G and Lp have been reported previously (Zheng et al., 2025).

**Table 1.** Monkey information and task parameters

| Monkey | Electrodes | Hand | Target | Target initial | Delay | Sessions |
| --- | --- | --- | --- | --- | --- | --- |

|  |  |  | velocity (°/s) | locations | length (s) |  |
| --- | --- | --- | --- | --- | --- | --- |
| **B** | Two 96-ch Utah arrays | Right | 0, $\pm 81$, $\pm 162$ | 8 | 0.5–1 | 1 |
| **Lb** | Two 96-ch + one 64-ch Utah arrays | Left | 0, $\pm 120$, $\pm 240$ | 8 | 0.6–1 | 3 |
| **G** | 64-ch S-probe | Left | 0, -120, -240 | 4 | 0.2 or 1 | 12 |
| **Lp** | 64-ch S-probe | Right | 0, -120, -180 | 4 | 0.2 or 1 | 14 |

### 2.3. Neural signal extraction

#### 2.3.1. Spike band power (SBP)

To extract the spike band power (SBP), raw neural signals were first common average referenced (CAR) by subtracting the mean signal across all recording channels at each time point. For each channel, the CAR-referenced signal was band-pass filtered between 300 and 1000 Hz using a fourth-order Butterworth filter to isolate the spike frequency band. The filtered signal was subsequently downsampled from 30 kHz to 2 kHz using finite impulse response (FIR) decimation with zero-phase filtering to prevent phase distortion. The instantaneous signal magnitude was obtained by full-wave rectification (absolute value) followed by temporal smoothing using a moving-average filter with a window length of 100 samples (50 ms at 2 kHz). The smoothed SBP signal was finally downsampled to 1 kHz for subsequent analyses.

#### 2.3.2. Local motor potentials (LMP)

Broadband neural recordings sampled at 30 kHz were low-pass filtered at 1000 Hz using a second-order Butterworth filter and decimated to 2 kHz. A fourth-order Butterworth low-pass filter with a cutoff frequency of 500 Hz was subsequently applied to obtain LFP signals. The LMP was then computed by convolving each LFP signal with a normalized Hanning window of 50 ms. LMP power was computed by squaring the LMP signal and applying a 10-sample moving average.

#### 2.3.3. Local field potential mean power

Frequency-specific LFP power features were extracted from the preprocessed LFP signals. Five canonical frequency bands were analyzed: delta (0.5–4 Hz), beta (12–25 Hz), low gamma (50–100 Hz), high gamma (100–200 Hz), and ultra-high gamma (200–400 Hz). For each frequency band, the LFP signals were band-pass filtered using a third-order Butterworth filter with zero-phase forward and reverse filtering to avoid phase distortion. Band-specific power was estimated using the sliding-window approach. For each time window, the mean-squared amplitude of the filtered signal was computed. To account for inter-electrode variability, band power was normalized using z-score transformation relative to a predefined baseline period (MO-0.5s to MO-0.3s).

### 2.4. Spike sorting

Neural signals were offline high-pass filtered at 300 Hz (second-order Butterworth filter) and exported as 16-bit signed integer binary files per channel for spike sorting in Plexon Offline Sorter (OFS; Plexon Inc.). Spikes were detected using a negative threshold set at −4.5×RMS of the background noise. Waveforms were represented using principal component analysis (PCA), and initial spike clusters were identified using the Valley-Seeking algorithm. Clusters were then manually curated based on waveform morphology, cluster separation, and inter-spike interval distributions. Units with well-isolated, consistent waveforms and minimal refractory-period violations were retained. Final spike timestamps and unit assignments were exported in Plexon (.plx) format for further analysis.

### 2.5. Neural decoders and decoding process

To evaluate the movement relevance of different neural features, we decoded the two-dimensional hand position and velocity during movement execution, defined as the period from movement onset to the target touch, using neural features. For each recording session, the execution periods from individual trials were concatenated sequentially in their original temporal order. All 64 channels were used for the S-probe and 64-channel Utah array recordings, whereas 64 channels were randomly selected from the 96-channel Utah array to ensure a matched channel count across electrode configurations. For consistency across the recording sessions, the neural and kinematic data were truncated to the minimum session length across the dataset. Neural signals were sampled at 100 Hz for subsequent decoding analysis. For each neural feature, separate decoders were trained to predict the kinematics, with the exception of preferential subspace identification (PSID), where four kinematic parameters were decoded from the same fitted model. Decoding performance was quantified using the coefficient of determination (*R*^2^) between the predicted and actual kinematic variables. Unless otherwise specified, all decoders were trained and evaluated using identical neural features and movement variables under the same fivefold cross-validation framework. Specifically, the concatenated neural and kinematic data were divided into five contiguous temporal blocks without shuffling. In each fold, one block was held out for testing, while the remaining four blocks were used for model training. The hyperparameters for each decoder were optimized using an exhaustive grid search over predefined parameter ranges. For each candidate value, the decoding performance was independently evaluated across the recording sessions for two-dimensional position and velocity decoding. The decoding performance was quantified using *R*^2^ for both position and velocity decoding. The optimal parameter set was selected based on the highest mean *R*^2^ across the sessions and decoding targets (**Table 3**).

**Table 2.** PSID hyperparameters

| Features | Electrode | $n_x$ (median [95% CI]) | $n_1$ (median [95% CI]) |
| --- | --- | --- | --- |
| **LMP** | **Utah** | 20.0 [19.0 20.0] | 9.0 [6.0 17.0] |
|  | **S-probe** | 18.5 [13.0 20.0] | 5.0 [2.0 8.0] |
| **Power 0.5–4 Hz** | **Utah** | 15.0 [11.0 16.0] | 2.0 [2.0 4.0] |
|  | **S-probe** | 9.5 [5.0 18.0] | 2.0 [2.0 2.0] |
| **Power 12–25 Hz** | **Utah** | 10.0 [8.0 11.0] | 2.0 [2.0 3.0] |
|  | **S-probe** | 11.0 [8.0 13.0] | 2.0 [2.0 2.0] |
| **Power 50–100 Hz** | **Utah** | 11.0 [7.0 11.0] | 2.0 [2.0 3.0] |
|  | **S-probe** | 11.0 [11.0 12.0] | 2.0 [2.0 2.5] |
| **Power 100–200 Hz** | **Utah** | 15.0 [10.0 19.0] | 4.0 [3.0 5.0] |
|  | **S-probe** | 11.5 [10.5 16.0] | 3.0 [2.0 4.0] |
| **Power 200–400 Hz** | **Utah** | 14.0 [11.0 19.0] | 6.0 [4.0 13.0] |
|  | **S-probe** | 13.0 [11.0 16.5] | 5.0 [3.0 7.0] |
| **SBP** | **Utah** | 18.0 [16.0 20.0] | 15.0 [10.0 16.0] |
|  | **S-probe** | 17.0 [15.5 20.0] | 9.0 [7.0 11.0] |

CI: confidence intervals

**Table 3.**
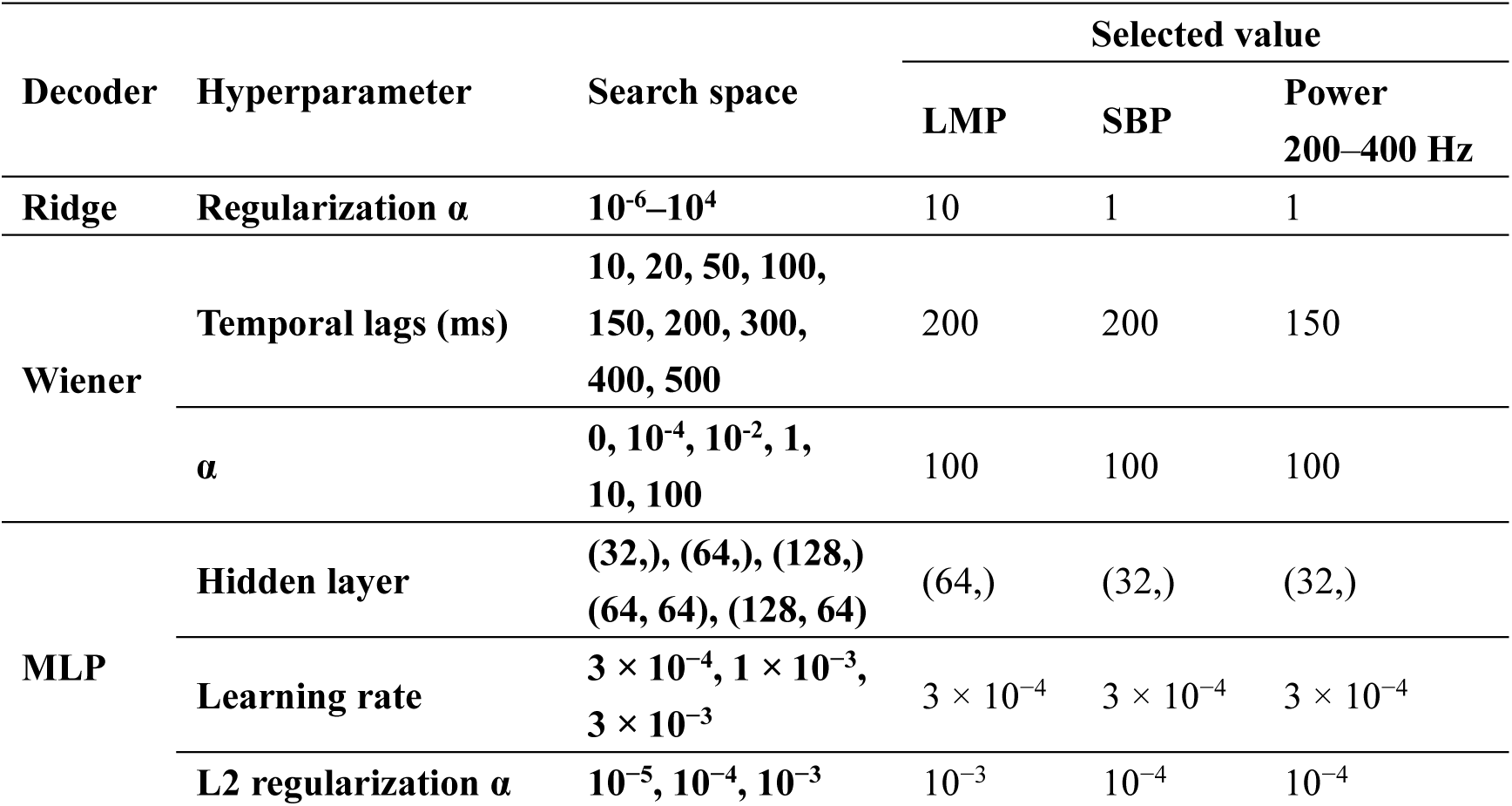
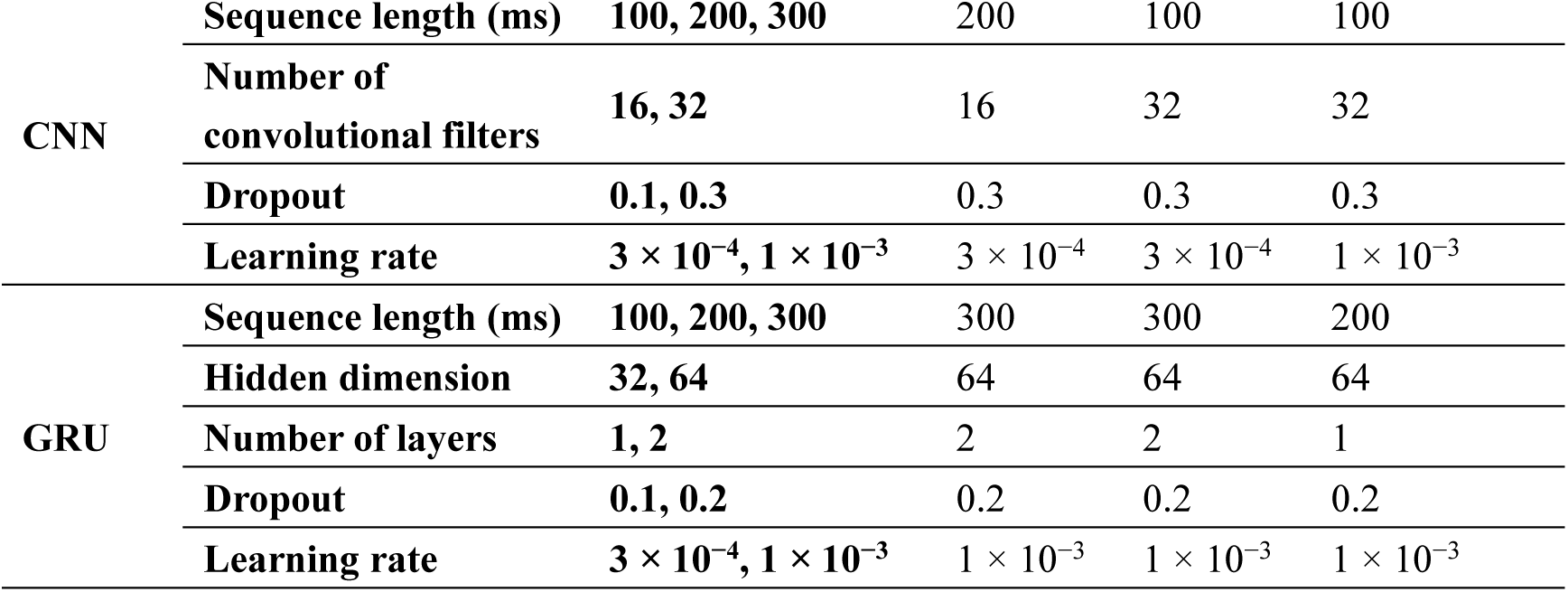
Decoder hyperparameters

#### 2.5.1. Ridge regression decoder

Ridge regression was introduced as a regularized linear baseline. Prior to model training, each neural feature was standardized to a zero mean and unit variance using statistics computed from the training set. Given the neural feature vector **x**(*t*) at time *t*, the decoder assumes a linear relationship with movement variables.

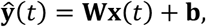

where **W** denotes the decoding weight matrix, and **b** is a bias term. The decoding weights are estimated by minimizing the regularized least-squares objective:

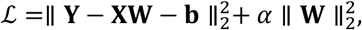

where **X** denotes the neural feature matrix, **Y** denotes the corresponding movement variable, and *α* denotes the L2 regularization coefficient. The L2 penalty reduces model variance and improves numerical stability in the presence of correlated neural features. The regularization coefficient (*α*) was optimized using an exhaustive grid search over 11 values spanning 10^-6^ to 10^4^.

#### 2.5.2. Wiener filter decoder

Compared with conventional ridge regression, the Wiener filter explicitly incorporates the temporal history of neural activity, allowing it to capture the temporal dependencies between neural signals and behavioral outputs. The behavioral output at time *t* was predicted as a weighted linear combination of neural activity over the preceding (L) time bins as follows:

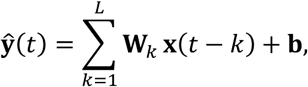

where **x**(*t* − *k*) denotes the neural feature vector recorded *k* time bins before the prediction time, **W***_k_* is the decoding weight matrix, and **b** is a bias term. The neural activity from the preceding (*L*) time bins was concatenated into a single feature vector, and the decoding weights were estimated using ridge regression. The hyperparameters of the Wiener filter were optimized using an exhaustive grid search over the number of temporal lags (*L*: 1, 2, 5, 10, 15, 20, 30, 40, and 50 samples, corresponding to temporal windows of 10–500 ms at the 100-Hz decoding rate) and ridge regularization coefficient (*α*: 0, 10^-4^, 10^-2^, 1, 10, and 100), resulting in 54 parameter combinations.

#### 2.5.3. Preferential Subspace Identification (PSID)

Conventional linear decoders do not explicitly separate behaviorally relevant neural dynamics from behaviorally irrelevant neural variabilities. To better capture the latent neural dynamics underlying behavior, we employed Preferential Subspace Identification (PSID), a linear state-space modeling framework that explicitly prioritizes latent states predictive of behavior while simultaneously modeling the remaining neural variance (Sani *et al*., 2021). Neural population activity (*y_k_* ∈ ℝ*^ny^*) and behavioral variables (*z_k_* ∈ ℝ*^nz^*) were modeled using a general dynamic linear state space system:

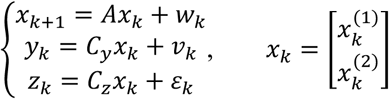

where k specifies the time index, *x_K_* ∈ ℝ*^nx^* denotes the latent state,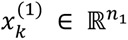 and 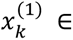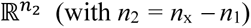 are its behaviorally relevant and irrelevant components, respectively. *ε_k_* represents behavior dynamics that are not present in the recorded neural activity, and *w_k_* and *v_k_* are zero-mean white noises. The matrices *A*, *C_y_*, and *C_z_* describe the latent dynamics and observation models. The total latent-state dimension (*n_x_*) and number of behaviorally relevant latent states (*n*_1_) were selected through a grid search. For every signal representation, the PSID models were trained over a grid of latent-state dimensions (*n_x_* = 2–20) and behaviorally relevant dimensions (*n*_1_ = 2, …, *n_x_*), with the prediction horizon fixed at *i* = 10. Model performance was evaluated on the held-out test set using the coefficient of determination (*R*^2^), and the combination of *n_x_* and *n*_1_ yielding the highest mean *R*^2^ across behavioral dimensions was selected as the optimal model for the recording session and feature type (**Table 2**). The identified model was subsequently used to predict behavioral variables from neural activity using the Kalman state estimation.

#### 2.5.4. Multilayer Perceptron (MLP) Decoder

A multilayer perceptron (MLP) was implemented to model the nonlinear relationships between neural activity and behavioral variables. Prior to training, each neural feature was standardized to a zero mean and unit variance using statistics computed from the training set. The network consists of customized hidden layers, followed by rectified linear unit (ReLU) activation functions. The output layer comprises two linear neurons corresponding to the two-dimensional kinematic variables. The network parameters were optimized using the Adam optimizer by minimizing the mean squared error (MSE) between the predicted and measured kinematic variables. MLP hyperparameters were optimized using an exhaustive grid search over a hidden layer architecture (single-hidden-layer networks with 32, 64, or 128 neurons and two hidden layer networks with 64 and 64 or 128 and 64 neurons), learning rate (3 × 10^−4^, 1 × 10^−3^, and 3 × 10^−3^), and L2 regularization strength (α = 10^−5^, 10^−4^, and 10^−3^), yielding 45 parameter combinations. For each combination, the MLP was trained and evaluated independently for each recording session using standardized neural features and early stopping (maximum 500 iterations; patience = 15).

#### 2.5.5. Multi-scale Temporal Convolutional Neural Network (CNN) Decoder

A multi-scale convolutional neural network (CNN) was employed to learn temporal patterns in neural activity at multiple time scales. The neural features were standardized and segmented into overlapping windows before being presented to the network. The network consisted of three parallel one-dimensional convolutional branches with kernel sizes of 3, 7, and 15 samples, enabling the extraction of short-, intermediate-, and long-timescale temporal features, respectively. The outputs of the three branches were concatenated and normalized using batch normalization, followed by a 1 × 1 convolution to integrate the information across the feature channels. The temporal information was subsequently aggregated using depthwise one-dimensional convolution with a kernel size of 5. The feature vector from the final time step was passed through a fully connected prediction head comprising a linear layer, ReLU activation, dropout, and final linear layer to estimate the two-dimensional kinematic variables. The network parameters were optimized using the Adam optimizer to minimize MSE loss. CNN hyperparameters were optimized using an exhaustive grid search over the temporal sequence length (10, 20, and 30 samples, corresponding to 100, 200, and 300 ms at a sampling rate of 100 Hz), learning rate (3 × 10^−4^ and 1 × 10^−3^), number of convolutional filters (16 and 32), and dropout rate (0.1 and 0.3), yielding 24 parameter combinations. The hidden dimensionality was fixed at 64 across all configurations. For each parameter combination, the CNN was trained and evaluated independently for each recording session, for both position and velocity decoding. Training was performed for 50 epochs using minibatches of 256 samples.

#### 2.5.6. Bidirectional gated recurrent unit (BiGRU) Decoder

To exploit the temporal dependencies in neural activity, recurrent neural network decoders based on bidirectional gated recurrent unit (BiGRU) architectures were implemented. Neural features were first standardized and segmented into overlapping temporal windows. The recurrent network consists of bidirectional recurrent layers, with the number of layers and hidden dimensionality optimized through a grid search. The forward and backward hidden states were concatenated to form a joint temporal representation. The hidden state at the final time step was passed through a fully connected prediction head comprising a linear layer, ReLU activation, dropout, and final linear layer to estimate the two-dimensional kinematic variables. The model parameters were optimized using the Adam optimizer to minimize MSE loss. Ten percent of the training sequences was reserved for validation. The learning rate reduction on plateau and early stopping (patience = 10 epochs) were applied to reduce overfitting, and the model with the lowest validation loss was retained for evaluation. BiGRU hyperparameters were optimized using an exhaustive grid search over the temporal sequence length (10, 20, and 30 samples, corresponding to 100, 200, and 300 ms at a sampling rate of 100 Hz), hidden dimensionality (32 and 64 units), number of recurrent layers (1 and 2), dropout rate (0.1 and 0.2), and learning rate (3 × 10^−4^ and 1 × 10^−3^), yielding 48 parameter combinations. For each combination, the BiGRU decoder was independently trained and evaluated for each recording session for both the position and velocity decoding. A batch size of 256 and maximum of 100 training epochs were used.

### 2.6. Spatial correlation analysis

To quantify the spatial organization of neural activity, pairwise Pearson correlation coefficients were computed between all the recording channels within each recording session. Prior to correlation analysis, neural signals from each channel were linearly detrended to remove the slow baseline drift while preserving the intrinsic correlation structure. For each electrode pair, the Euclidean distance between the recording sites was calculated using physical electrode geometry. The pairwise correlation coefficients were grouped according to their inter-electrode distances.

For the Utah arrays, all the channel pairs were included in the analysis. For linear probes, electrode pairs were classified into vertical pairs (channels located within the same probe column) and cross pairs (channels located in different columns), allowing spatial correlations along and across the probe to be analyzed separately. For population-level analysis, distance-dependent correlation curves from individual recording sessions were pooled using inter-electrode distance. At each distance, the median correlation coefficient across sessions was calculated and 95% confidence intervals (CIs) were estimated by bootstrap resampling (100 resamples). The pooled median correlation profile was subsequently fitted using an exponential decay model to estimate the group-level spatial correlation length.

### 2.7. Preferred direction estimation

The preferred directions (PDs) were estimated for each recording channel using a cosine-tuning model based on the direction-dependent neural responses. For each movement direction, neural activity was first baseline-corrected by subtracting the mean activity during the baseline period (−300 to 0 ms relative to movement onset) from the mean activity during the task period (0–300 ms), yielding a direction-specific response. The direction-tuning curve was fitted using a first-order cosine model:

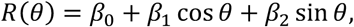

where *β*_0_ represents the direction-independent intercept, and *β*_1_ and *β*_2_ describe the directional modulation. The preferred direction is defined as

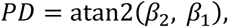

and the tuning amplitude was calculated as

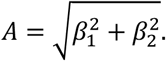

The goodness of fit was quantified using the coefficient of determination (*R*^2^).

The PD reliability was assessed using bootstrap resampling and permutation testing. For 1000 bootstrap iterations, trials were resampled with replacement within each movement direction, and the cosine model was refitted to estimate the PD and tuning amplitude. Circular 95% CIs for PD and 95% CIs for tuning amplitude were derived from the corresponding bootstrap distributions. Directional tuning significance was assessed using 1000 trial-level permutations by shuffling movement-direction labels while preserving neural responses. The permutation p-value was calculated as the proportion of permuted tuning amplitudes that were greater than or equal to the observed amplitude. A recording channel was considered directionally tuned only if it simultaneously satisfied three criteria: (1) the bootstrap circular confidence interval width was smaller than *π* radians, indicating a well-defined PD; (2) the permutation test yielded *p* < 0.05; and (3) the cosine model explained a sufficient proportion of the response variance (*R^2^* > 0.2).

Only channels meeting all three criteria were included in the subsequent population analyses.

### 2.8. Mean resultant length of preferred directions

The mean resultant length (*R*) was calculated to quantify the concentration of PDs across neural channels for each recording session. Each PD is represented as a unit vector in the circular domain and *R* is computed as the magnitude of the mean resultant vector:

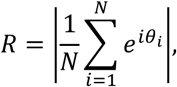

where *N* is the number of channels, and *θ_i_* is the PD of the *i*-th channel. *R* ranges from 0 to 1, with higher values indicating a stronger clustering of PDs and lower values indicating a greater dispersion of PDs across channels.

### 2.9. Channel importance and cumulative decoding performance

To characterize the relationship between the number of included channels and decoding performance, we first quantified the contribution of individual neural channels to decoding performance by performing a leave-one-channel-out analysis using the PSID decoder. The PSID model was fitted with a latent state dimensionality of (*n_x_* = 10), task-related state dimensionality of (*n*_1_ = 5), and integration order of (i = 10). For each fold, a baseline PSID model was first trained using all available channels and the decoding performance was quantified using the coefficient of determination (*R^2^*) between the predicted and actual kinematic variables. To estimate the contribution of each channel, the channel was individually removed from the neural feature set, the PSID model was refitted using the remaining channels, and the decoding performance was recalculated for the same held-out test segment. Channel importance is defined as the reduction in decoding performance resulting from channel removal.

Next, channels were ranked according to their mean dropout-derived importance and then progressively included according to this ranking, and the PSID models were refitted using the top 1%, 2%, 5%, 10%, 20%, 30%, 40%, 50%, 75%, and 100% of channels, rounded to the nearest integer number of channels. For each channel subset, the decoding performance was evaluated on the corresponding held-out test segments across all the cross-validation folds. Therefore, the resulting decoding-performance curves quantify how rapidly the decoding performance approaches its maximum as the most informative channels are progressively incorporated. For group-level analysis, the session-level decoding performance at each channel proportion was summarized as the mean ± standard error of the mean (SEM) across recording sessions.

Additionally, the channel proportion required to reach 95% of the maximum decoding performance was determined for each group as the smallest channel proportion at which the decoding performance exceeded 95% of the group’s maximum observed performance.

### 2.10. Frequency-band feature extraction

To evaluate the decoding performance of the raw signals and band-limited power of different temporal components of the LMP, we extracted distinct frequency bands from the LMP: delta (0.1–4 Hz), theta (4–8 Hz), alpha (8–13 Hz), and beta (13–20 Hz). For each frequency band, two feature representations were generated: (1) the band-limited raw signal, obtained by fourth-order Butterworth band-pass filtering, and (2) the corresponding band power, computed as the squared amplitude, followed by temporal smoothing using a moving average filter (window length = 10 samples). To further characterize the frequency dependence of the decoding performance, a series of low-pass filtered signals from the LMP was generated using cutoff frequencies of 1, 2, 3, 4, 5, 8, 10, 15, 20, 30, and 40 Hz. All filters were implemented as fourth-order zero-phase Butterworth filters by using forward and reverse filtering.

### 2.11. Movement-related cortical potentials (MRCP) analysis

To characterize movement-related cortical potentials (MRCPs), MRCP signals were obtained by band-pass filtering of continuous LMP signals between 0.1 and 4 Hz using a fourth-order zero-phase IIR filter. Neural activity was then segmented into epochs ranging from 1 s before to 0.5 s after movement onset (MO). Trials were grouped according to the target speed. For each speed condition, a representative response was first obtained for each trial by computing the median waveform across the recording channels. The population responses were then estimated as medians across the trials. To quantify variability, 95% CIs were calculated using bootstrap resampling of trials (1,000 bootstrap iterations). To determine whether movement-related activity differed across target speeds, each non-zero-speed condition was compared with the stationary condition (0°/s). Statistical significance at each time point was assessed using a non-parametric permutation test (2,000 permutations), in which trial labels were randomly shuffled between two groups while preserving group sizes. The observed difference in the median responses was compared with the permutation distribution to obtain two-sided *p*-values. Multiple comparisons across time were controlled using the Benjamini–Hochberg false discovery rate (FDR) procedure with a significance threshold of 0.05. The time points that survived the FDR correction are displayed beneath the corresponding waveform.

## 3. Results

### 3.1. Behavioral and neural data preprocessing

To compare the neural features across the two electrode configurations, we analyzed datasets from our previous studies in which monkeys performed a manual delayed interception task. In this task, peripheral targets were either static (classic center-out reach) or moving (interception). Because the reach trajectories during movement execution were highly similar between the two conditions (Li, Wang and Cui, 2017; Zheng, Wang and Cui, 2025), we treated the two conditions together in most subsequent analyses (**figure 1(a)**). From one study, monkey B contributed one recording session recorded with two 96-channel Utah arrays targeting the primary motor cortex (M1) and dorsal premotor cortex (PMd). Monkey Lb contributed three recording sessions, with two 96-channel Utah arrays targeting M1 and PMd, and one 64-channel array targeting the dorsolateral prefrontal cortex (dlPFC). Recordings from each cortical region were treated as an independent dataset, with neural signals and behavioral data from the corresponding recording site analyzed separately. Thus, each recording session contributed two region-specific datasets for monkey B and three region-specific datasets for monkey Lb. In total, 11 datasets were obtained from recordings with Utah arrays (**figure 1(b)**). From another study, Monkeys G and Lp contributed 12 and 14 datasets, respectively, with acute recordings from either M1 or PMd using 64-channel two-column linear S-probes (Zheng, Wang and Cui, 2025).

**Figure 1.**
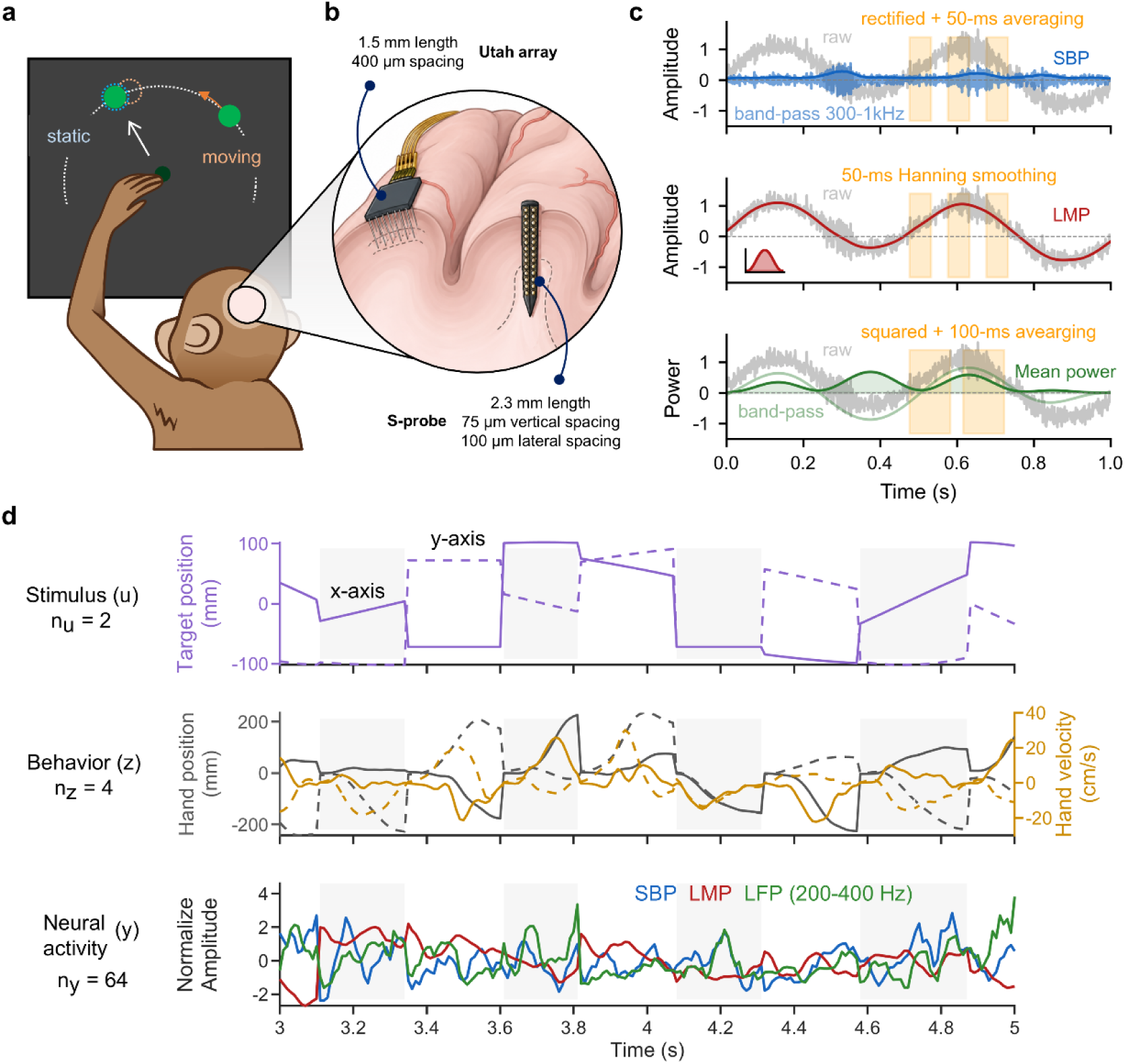
Experimental paradigm, recording configuration, signal processing. **(a)** Schematic of the delayed manual interception task. Monkeys initiated reaches toward either a stationary target (static) or a moving target (moving). **(b)** Neural recording configuration. Recordings were obtained using either a 64-channel linear probe (S-probe) or a 96- or 64-channel Utah microelectrode array implanted in the sensorimotor cortex. **(c)** Extraction of representative neural features, spike-band power (SBP), local motor potential (LMP) and mean power of local field potentials (LFP) from broadband signals. Shaded regions illustrate the temporal averaging windows used for feature extraction. **(d)** Behavioral and neural data for analysis. The stimulus vector (u) consisted of the two-dimensional target position. The behavioral state (z) comprised two-dimensional hand position and velocity. The neural observation (y) consisted of simultaneously recorded neural features from all recording channels. Alternating gray-shaded and blank regions distinguish individual trials.

We extracted three typical types of broadband neural signals: spike band power (SBP) (Nason *et al*., 2020), local motor potentials (LMP) (Schalk *et al*., 2007; Stavisky *et al*., 2015), and local field potential (LFP) mean power of different frequency bands including delta (0.5–4 Hz), beta (12–25 Hz), low gamma (50–100 Hz), high gamma (100–200 Hz), and ultra-high gamma (200– 400 Hz) (Gallego-Carracedo *et al*., 2022). SBP was computed as the power of high-frequency neural activity in the spike band, providing a population-level measure of the multiunit spiking activity. The LMP was obtained by low-pass filtering of the broadband neural signal, which primarily reflects low-frequency population activity. The LFP mean power was calculated as the average spectral power within predefined frequency bands of the LFP, providing a frequency-specific measure of oscillatory neural activity (**figure 1(c)**). As SBP, LMP, and high-gamma bands have been shown to exhibit strong correlations with movement kinematics, it is of interest to determine whether their decoding performance remains consistent across different electrode configurations.

Movement kinematics, including hand position and velocity, were obtained from the hand position tracked using an optical motion capture system. Neural activity was downsampled to 100 Hz to match the temporal resolution of the movement parameters. For each recording session, both movement and neural data corresponding to the execution periods (from movement onset to target touch) of individual trials were concatenated sequentially for subsequent analysis (**figure 1(d)**).

### 3.2. Decoding accuracy across neural modalities

First, we assessed the decoding performance of different signal modalities. Consistent with previous studies, the LMP and SBP achieved the highest *R^2^* for both hand positions, whereas the LFP mean power showed progressively higher *R^2^* at higher frequencies (**figure 2(a)**). Notably, the Utah-array sessions exhibited significantly higher decoding performance than the S-probe sessions, with a more pronounced difference between the two electrode configurations for LMP (LMP: median 0.51 [95% CI, 0.445–0.589] vs. 0.11 [0.038, 0.137]; mean power (200–400 Hz): 0.18 [0.131, 0.252] vs. 0.08 [0.028, 0.121]; SBP: 0.50 [0.480, 0.581] vs. 0.42 [0.349, 0.462]). The results were similar for hand-velocity decoding (**figure 2(b)**, LMP: 0.50 [0.465, 0.603] vs. 0.04 [0.005, 0.069]; mean power (200–400 Hz): 0.15 [0.120, 0.224] vs. 0.05 [0.027, 0.052]; SBP: 0.53 [0.488, 0.588] vs. 0.27 [0.198, 0.323]).

**Figure 2.**
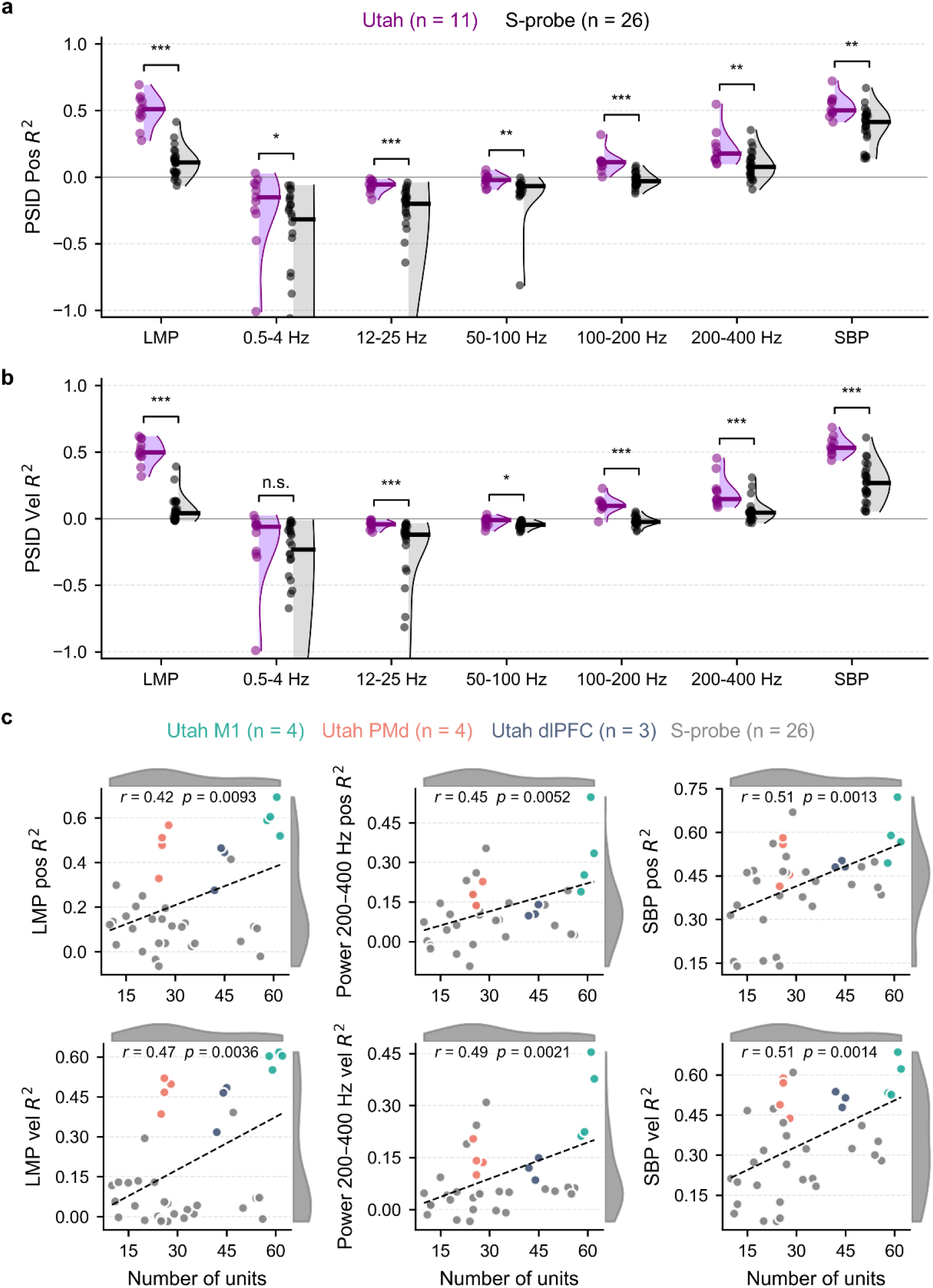
Decoding performance of different neural features and its relationship to neuronal population size. **(a)** Preferential subspace identification (PSID) decoding performance for hand position across different neural features, including LMP, SBP, and mean power in five frequency bands (0.5–4, 12–25, 50–100, 100–200 and 200–400 Hz). Data are grouped by electrode configurations. Each dot represents an individual recording session, with violin plots showing the distribution and horizontal bars indicating the median. **(b)** PSID decoding performance for hand velocity across different neural features. **(c)** Relationship between the number of recorded units and decoding performance for LMP, 200–400 Hz LFP mean power, and SBP. Each dot represents a recording session. Dashed lines indicate linear regression fits, with Pearson correlation coefficients (*r*) and *P* values shown in each panel.

Given the strong behavioral relevance of local spiking activity, the presence of single-unit activity in a recording channel may substantially influence the decoding performance of broadband neural signals. Therefore, we further examined the correlation between the single-unit count and *R^2^*. As expected, both LFP mean power and SBP showed a significant positive correlation with single-unit number across electrode configurations, suggesting that the difference observed between Utah-array sessions and S-probe sessions of these two neural modalities was mainly attributed to the relatively fewer single-units recorded in S-probe sessions (**figure 2(c)**).

However, for LMP, the Utah-array sessions deviated from the S-probe sessions, exhibiting a much higher *R^2^* than the S-probe sessions with similar single-unit counts. In line with this observation, the distribution of LMP *R^2^* values appeared to exhibit a bimodal pattern, with the Utah-array and S-probe sessions forming two partially separated distributions.

These results suggest that although both LMP and SBP exhibited strong decoding performance, LMP recorded with the Utah array showed higher behavioral relevance than that recorded with the S-probe. Notably, this advantage was not explained by the presence of single-unit activity, which is consistent with the low frequency of LMP and its relatively limited influence from high-frequency spiking activity. Therefore, we next investigated the factors underlying the difference in decoding performance between the two electrode configurations.

### 3.3. Spatial organization of neural signals across recording electrodes

Given that behavioral dimensionality and encoding capacity are related to neural heterogeneity across channels, we first compared the across-channel correlations across different neural modalities and electrode configurations. The mean correlation results revealed that LMP exhibited the highest across-channel correlations, whereas SBP exhibited the lowest. The LFP mean power showed intermediate levels of cross-channel correlation, which generally decreased with increasing frequency, with the exception that the 12–25 Hz band, rather than the 0.5–4 Hz band, showed the lowest correlation among the LFP frequency bands (**figure 3(a)**, LMP: 0.97 [0.533, 0.989]; 0.5–4 Hz: 0.42 [0.162, 0.688]; 12–25 Hz: 0.78 [0.223, 0.972]; 50–100 Hz: 0.45 [0.126, 0.89]; 100–200 Hz: 0.21 [0.055, 0.856]; 200–400 Hz: 0.11 [0.016, 0.843]; SBP: 0.089 [0.026, 0.343]). As expected, Utah-array sessions exhibited a lower correlation than S-probe sessions, especially for LMP and the 12–25 Hz frequency band. In addition, Utah array recordings from different brain areas showed no substantial differences in cross-channel correlations.

**Figure 3.**
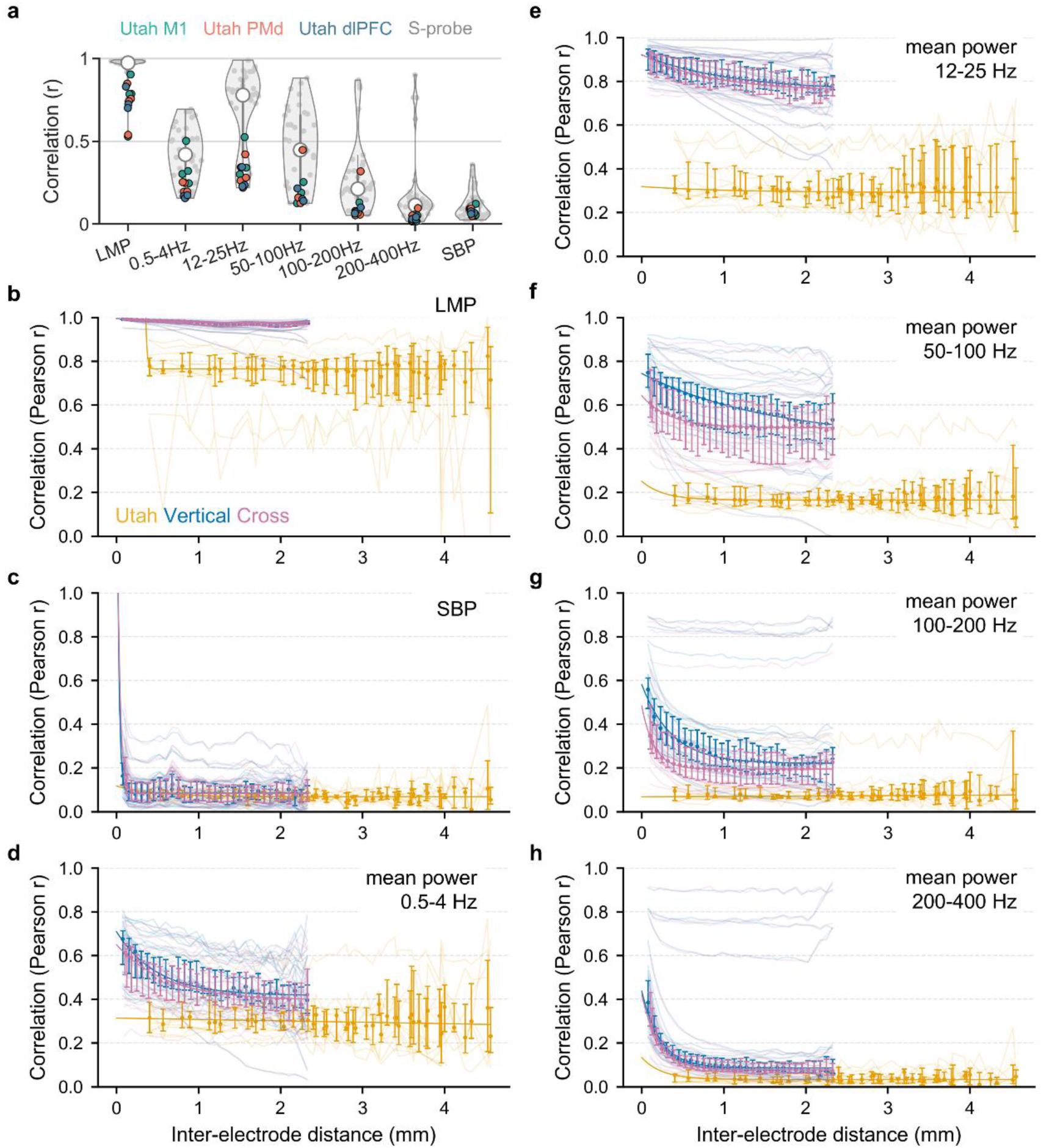
Spatial organization of neural signals across recording electrodes. **(a)** Distribution of pairwise Pearson correlations as a function of neural signal type across Utah arrays and S-probes. (b–h) Pearson correlation between electrode signals as a function of inter-electrode distance for LMP **(b)**, SBP **(c)**, and mean power in the 0.5–4 Hz **(d)**, 12–25 Hz **(e)**, 50–100 Hz **(f)**, 100–200 Hz **(g)** and 200–400 Hz **(h)** bands. “Vertical” indicates channel pairs located within the same probe column; “Cross” denotes channel pairs located across different probe columns. Thin lines represent individual recording sessions, whereas solid lines with error bars represent the mean correlation ± s.e.m. across sessions at each inter-electrode distance.

As the Utah array and S-probe differed in electrode spacing and spatial extent, we further evaluated how cross-channel correlations varied as a function of inter-electrode distance. For the S-probe, electrode pairs were further classified according to their spatial arrangement: pairs spanning the two probe columns with the two electrodes horizontally aligned were defined as cross-column pairs, whereas pairs within the same column with vertically aligned electrodes were defined as vertical pairs. Consistently, LMP in S-probe sessions showed higher across-channel correlation than in Utah-array sessions at all inter-electrode distances, even when the distances overlapped with the Utah array (**figure 3(b)**). In contrast, SBP showed a similar correlation at the same inter-electrode distance between the two electrode configurations (**figure 3(c)**). The 0.5–4 Hz band showed an intermediate pattern between the LMP and SBP. In S-probe sessions, across-channel correlations decreased as the inter-electrode distance increased and progressively approached those observed in Utah-array sessions, suggesting that its channel correlation is primarily determined by the inter-electrode distance rather than the electrode configuration (**figure 3(d)**). For the other frequency bands, the correlations in the S-probe sessions generally decreased exponentially with increasing inter-electrode distance. Moreover, the difference between the S-probe and Utah-array sessions at comparable inter-electrode distances progressively diminished with increasing frequency (**figure 3 (e-h)**). Notably, in the 50–100 Hz and 100–200 Hz band, cross-column pairs showed a lower correlation than vertical pairs at comparable distances, supporting the idea that the information encoded by these frequency components may be more heterogeneous along the horizontal than the vertical cortical dimension.

These results suggest that for high-frequency signal modalities, such as SBP and the 200–400 Hz band, Utah-array and S-probe recordings exhibit similar channel correlations at comparable inter-electrode distances. In contrast, for low-frequency signals, particularly LMP, channels distributed along the vertical dimension exhibit more correlated information than those distributed along the horizontal dimension. This difference in the spatial organization of neural information may partially explain the higher decoding performance observed with Utah arrays.

### 3.4. Preferred directions distributions of neural signals

To further characterize the movement-related information encoded across different electrode configurations, we analyzed the preferred direction (PD) of the individual channels. According to the population vector theory, a broader distribution of PDs across channels provides more diverse directional information and may support robust population-level encoding (Georgopoulos *et al*., 1982; Georgopoulos, Schwartz and Kettner, 1986). Therefore, we examined whether Utah-array recordings exhibited greater PD diversity than S-probe recordings, potentially contributing to their superior decoding performances.

To test this hypothesis, we first averaged the neural activity across trials for each reach direction and aligned them to movement onset (MO). All signal modalities showed direction-dependent tuning, with distinct temporal activity profiles across the modalities (**figure 4(a)**). As expected, the tuning profiles of the frequency bands above 100 Hz increasingly resembled those of SBP, suggesting the contribution of similar underlying neural activity. Next, we computed the PDs based on neural activity from MO to MO+300 ms. Consistent with the peri-event time histogram (PETH) results, PDs estimated from high-frequency signal modalities were more similar to one another, whereas those derived from low-frequency components, including the LMP and the 0.5–4 Hz band, differed substantially (**figure 4(b)**). Despite some degree of depth-dependent variation, LMP PDs in the S-probe sessions were highly similar across channels, both within and across recording sessions from the same monkey. For example, LMP PDs were predominantly oriented toward approximately 4 o’clock in monkey G and 9 o’clock in monkey Lp (**figure 4(c)**). In the Utah array recordings, PDs were also highly similar across channels within each array, although they exhibited greater variability than those observed with the S-probe. In addition, Utah arrays recorded from different brain areas exhibited distinct PD distributions (**figure 4(d), left**).

**Figure 4.**
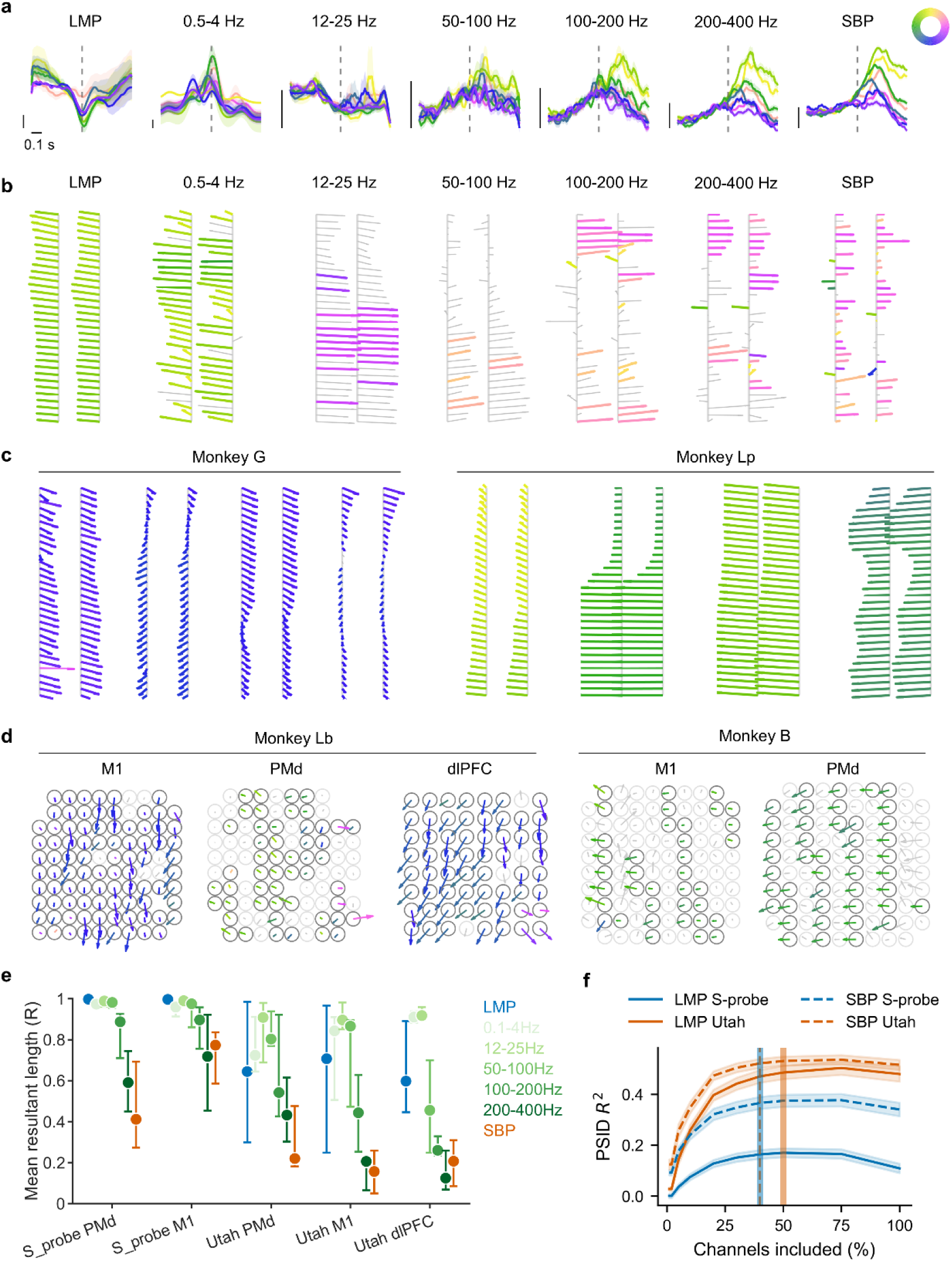
Preferred directions of neural features across electrode configurations. **(a)** Population-averaged neural responses in a representative Utah array recording session of monkey B. Curves indicate the mean responses and shaded areas indicate the s.e.m.; dashed vertical lines denote movement onset. The vertical scale represents 100 μV for LMP, 2 a.u. for LFP mean power, and 2 μV for SBP. **(b)** Preferred direction (PD) maps of a representative S-probe recording session of monkey Lp. Vector orientation and color encode PD angles; vector length is proportional to the normalized tuning amplitude. Electrodes with significant directional tuning are shown in color, whereas non-significant electrodes are shown in gray. **(c)** PD maps of multiple representative S-probe recording sessions in monkeys G and Lp. **(d)** PD maps from representative Utah array recording sessions in monkeys Lb and B across M1, PMd, and dlPFC. **(e)** Mean resultant length (*R*) of PDs for different electrode configurations and neural features. Points indicate the median and error bars indicate 95% confidence intervals. **(f)** Cumulative decoding analysis. Channels were ranked by their mean dropout-derived importance and progressively included according to this ranking. PSID decoding performance was summarized across cross- validation folds and recording sessions as mean ± SEM. Vertical color-coded lines indicate the minimum channel proportion required to reach 95% of the maximum decoding performance; solid lines are thicker than dashed lines for visual clarity.

Moreover, although channels with reliable PD estimates tended to show highly similar PDs, channels with less reliable PDs exhibited greater variability, potentially contributing to the more heterogeneous information encoded in the Utah-array recordings (**figure 4(d), right**).

Next, we quantified the PD concentration across the channels using the mean resultant length (*R*), defined as the magnitude of the mean resultant vector when each PD was represented as a unit vector. *R* ranges from 0 to 1, with higher values indicating more concentrated PDs and lower values indicating greater PD diversity across the channels. In the S-probe sessions, the *R* values of the LMP, 0.5–4 Hz, 12–25 Hz, and 50–100 Hz band were close to 1, indicating highly concentrated PD distributions. In contrast, *R* decreased progressively for the 100–200 Hz and 200–400 Hz bands and SBP (**figure 4(e)**). No significant differences were observed across the brain areas during the S-probe sessions. In the Utah-array sessions, although the relative pattern of *R* across signal modalities was similar to that in the S-probe sessions, *R* values were generally higher, particularly for LMP. Intriguingly, the *R* in the Utah-array sessions tended to be lower in the M1 and dlPFC than in the PMd. Therefore, these results indicate that LMP PDs were more variable in the Utah array than in the S-probe recordings. Consequently, when channels were sequentially included according to their dropout-derived importance ranking, LMP in S-probe sessions required a smaller proportion of channels to reach 95% of their maximum decoding performance than in Utah-array sessions (**figure 4(f)**, 40% vs. 50%). Notably, including additional channels in S-probe sessions could even reduce the LMP decoding performance, suggesting that highly correlated channels may provide redundant information and potentially introduce noise. In contrast, SBP required similar proportions (40%) of channels to approach a plateau decoding performance in the S-probe and Utah-array sessions.

In summary, broadband neural signals, particularly the LMP, exhibited distinct behavioral relevance across the Utah-array and S-probe recordings. This difference may be partially attributed to the greater heterogeneity of the information represented along the horizontal than the vertical cortical dimension, potentially contributing to the superior decoding performance of Utah-array recordings.

### 3.5. Spectral and temporal components underlying LMP decoding

The above analyses provided substantial evidence that the Utah-array recordings achieved a higher LMP decoding performance than the S-probe recordings. Given its high behavioral relevance and pronounced dependence on the electrode configuration, we next investigated the biophysical origin of LMP. Since LMP was first introduced by Schalk et al. (2007) and subsequently investigated in several studies, its underlying biophysical nature has remained unclear (Ahmadi, Constandinou and Bouganis, 2021). To date, the LMP has been primarily characterized as a low-frequency broadband signal with substantial value for neural decoding. Therefore, we first decomposed the LMP into four canonical frequency bands—delta (0.1–4 Hz), theta (4–8 Hz), alpha (8–13 Hz), and beta (13–20 Hz)—and evaluated their decoding performance using both the raw signals and their power. Notably, the raw delta-band signal achieved a decoding performance comparable to that of raw LMP, whereas the delta-band power yielded substantially lower *R^2^* values. In contrast, the raw signals of the other frequency bands showed consistently poor decoding performance, although their power yielded higher *R^2^*values in some Utah array recordings (**figure 5(a)**). These results suggest that the delta band contributes the most to the decoding performance of LMP.

**Figure 5.**
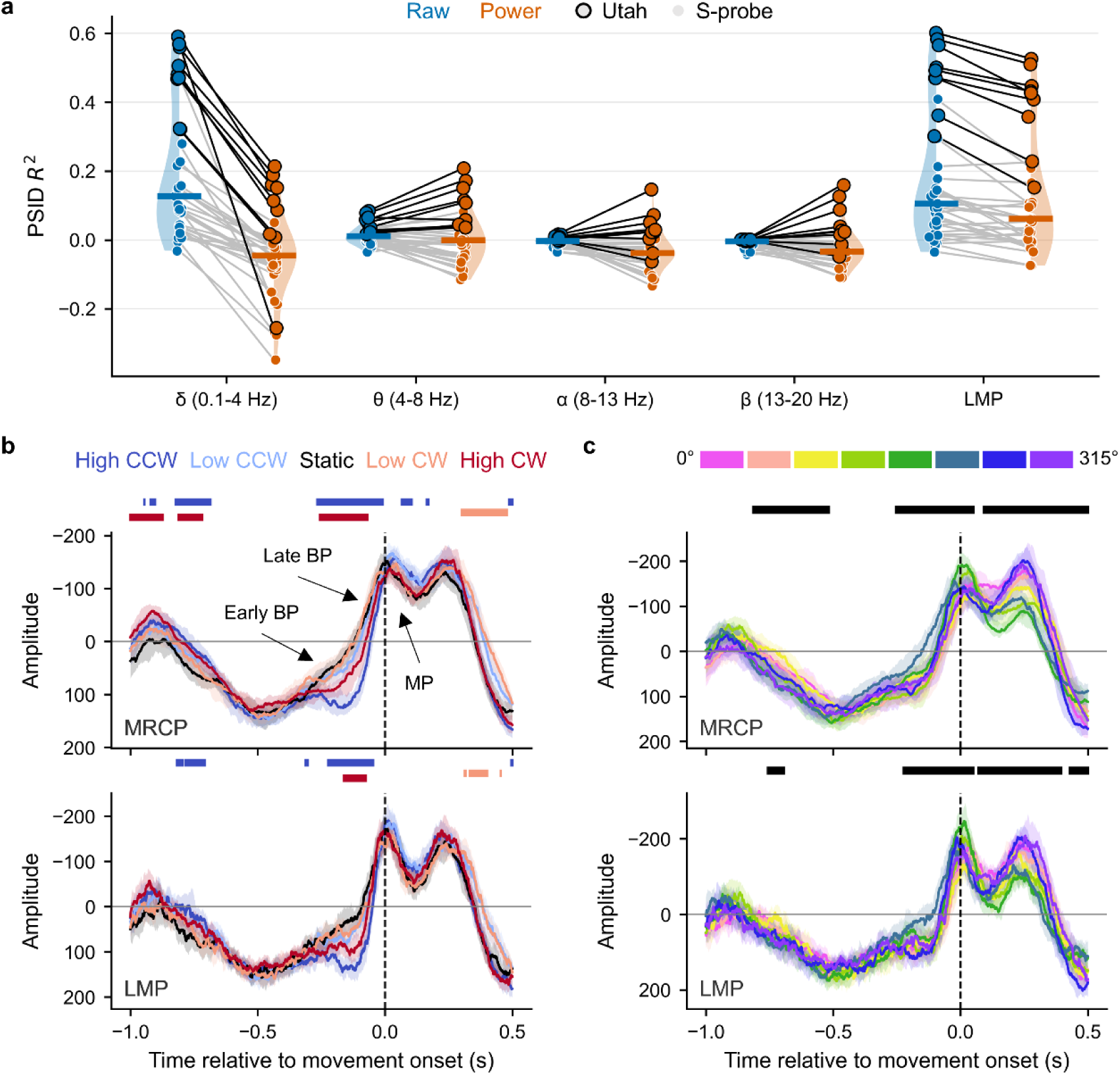
Decoding performance of different LMP frequency components and their correlation with movement-related cortical potentials (MRCPs). (a) PSID *R*^2^ values for raw LMP signals and power features across different frequency bands extracted from LMP. Each dot represents a recording session. Utah array recording sessions are highlighted with black edges. (b) Temporal evolution of MRCPs and LMP amplitudes relative to movement onset. Traces represent median values of trials grouped by target velocities. Shaded areas indicate 95% CIs, and horizontal bars indicate significant differences between each non-zero target velocity and the static condition. (c) Temporal evolution of MRCPs and LMP amplitudes with trials grouped by movement direction. Colors indicate eight movement directions, spaced at 45° intervals from 0° to 315° in the counterclockwise direction. Horizontal bars indicate time points at which the distributions differed significantly across directions, assessed using a time-resolved permutation-based Kruskal–Wallis omnibus test with false discovery rate correction for multiple comparisons.

The delta band range overlaps with a well-established neural signal, movement-related cortical potentials (MRCPs), which are slow cortical potentials associated with the preparation and execution of voluntary movement (Shibasaki and Hallett, 2006; Di Russo *et al*., 2017; Antenucci *et al*., 2026). To further examine the relationship between LMP and MRCP, we extracted the 0.1–4 Hz component from the neural signals as an MRCP-related signal and aligned both this component and LMP to MO. Notably, the LMP exhibited temporal characteristics close to those of MRCPs, with a gradual premovement buildup reminiscent of the early Bereitschaftspotential (BP), followed by an accelerated negative-going potential approaching the MO, consistent with the late BP, and a prominent motor potential (MP) around the MO. Both signals also differed across the target velocity conditions (**figure 5(b)**) and reach-direction conditions (**figure 5(c)**). Although MRCPs have been primarily defined and investigated using scalp or cortical surface-recorded EEG, MRCP-like intracortical activity may reflect related underlying neural mechanisms, albeit at a more local level.

Taken together, the overlap in the frequency range between the delta band and MRCPs suggests that the high behavioral relevance of the LMP is primarily attributable to its low-frequency components. However, the LMP power yielded higher *R^2^* values than delta-band power suggests that the LMP may also contain contributions from higher-frequency components. As these decoding results were obtained using a linear decoder, we hypothesized that the mixed spectral composition of the LMP may provide additional decoding advantages when nonlinear relationships are exploited by nonlinear decoders. In addition, although LMP has been investigated across various motor tasks (Ince *et al*., 2010; Flint *et al*., 2017), its decoding performance varies. One possible explanation is that the composite nature of the LMP may also introduce undesired noise that interferes with behaviorally relevant signals under certain conditions. Therefore, we further investigated these two hypotheses.

### 3.6. Decoding low-frequency LMP components across decoder architectures

To test the above hypotheses regarding LMP, we mainly used Utah-array sessions for subsequent analyses as they provided more stable and robust results. We first extracted signal components from the LMP using a series of low-pass cutoff frequencies (1, 2, 3, 4, 5, 8, 10, 15, 20, 30, and 40 Hz) and evaluated their decoding performance using both linear and nonlinear decoders. Linear decoders included ridge regression, Wiener filter, and PSID, whereas nonlinear decoders included multilayer perceptron (MLP), convolutional neural network (CNN), and gated recurrent unit (GRU). As expected, *R^2^* values generally increased with increasing cutoff frequency and reached a plateau after a cutoff frequency of 4 Hz for most decoders (**figure 6(a)**). At the plateau, GRU achieved the highest position decoding performance, followed by CNN, MLP, Wiener filter, PSID, and ridge regression (GRU: 0.75 ± 0.047; CNN: 0.71 ± 0.045; MLP: 0.55 ± 0.041; Wiener filter: 0.54 ± 0.037; PSID: 0.51 ± 0.036; ridge regression: 0.42 ± 0.038). For velocity decoding, the decoding performance ranked from highest to lowest as GRU, CNN, Wiener filter, MLP, PSID, and ridge regression (GRU: 0.68 ± 0.042; CNN: 0.67 ± 0.036; Wiener filter: 0.54 ± 0.034; MLP: 0.52 ± 0.032; PSID: 0.52 ± 0.032; ridge regression: 0.45 ± 0.029). Therefore, nonlinear decoders generally outperform linear decoders for LMP decoding.

**Figure 6.**
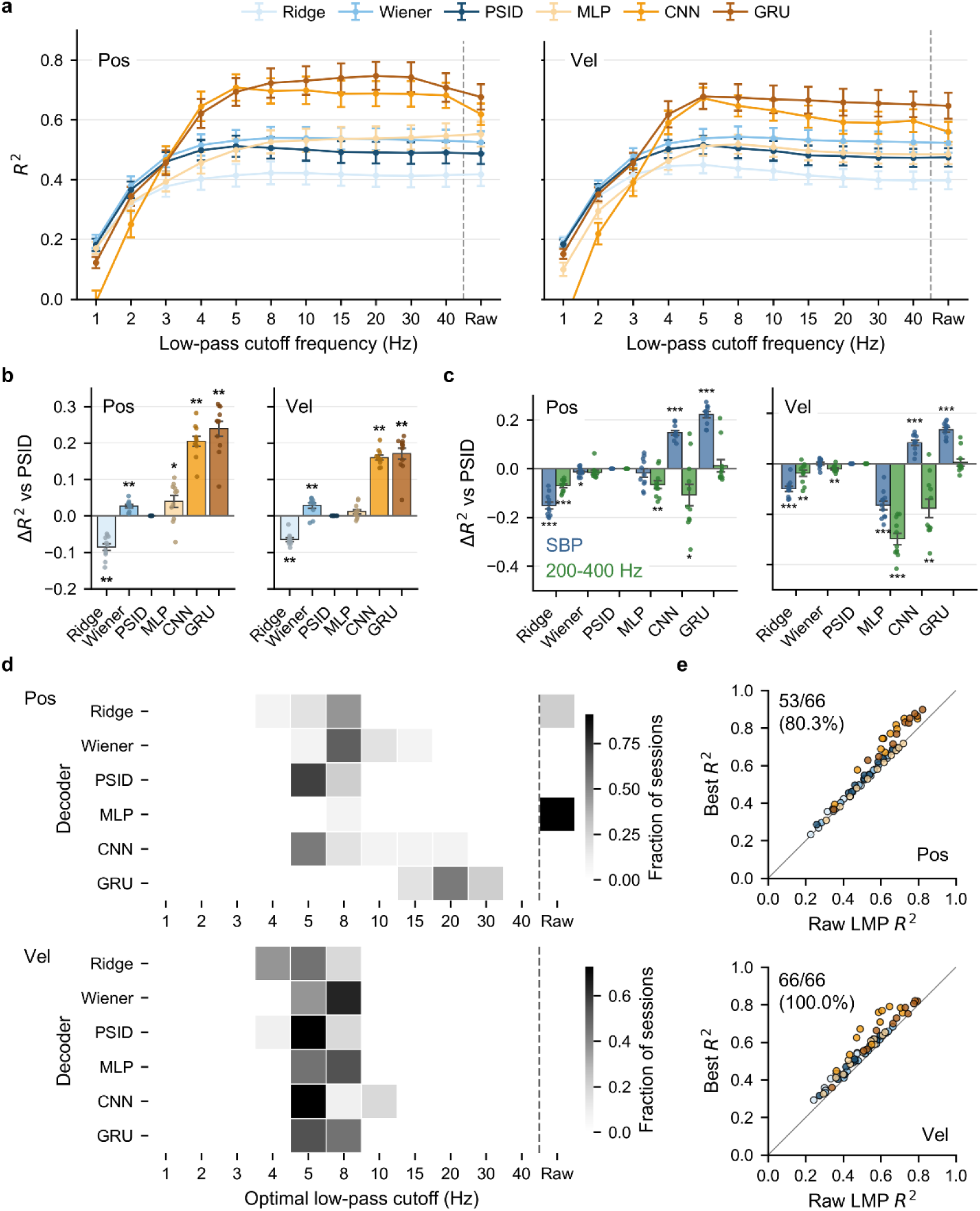
Evaluation of decoder performance and feature optimization in Utah-array sessions. **(a)** Position (Pos) and velocity (Vel) *R*^2^ across low-pass cutoff frequency for LMP-derived components and raw LMP signals using six decoders. Error bars denote standard error of the mean (s.e.m.) across sessions. Dashed lines separate extracted features from raw LMP for visual clarity. N = 11. **(b, c)** Relative decoding performance (Δ*R*^2^) compared to PSID across decoders for position and velocity using LMP features **(b)**, or SBP and 200-400-Hz band mean power **(c)**. Statistical comparisons between other decoders and PSID were performed using paired Wilcoxon signed-rank test with \*\**p* < 0.01, \*\*\**p* < 0.001. **(d)** Distributions of optimal low-pass cutoff frequencies across sessions for each decoder. **(e)** Comparison of the best *R*^2^ with *R*^2^ of raw LMP using each decoder. Dots are individual sessions color-coded by decoder types, as in panel (a). The proportion of sessions where the best *R*^2^ exceeded the raw LMP *R*^2^ is annotated.

To further quantify these performance differences, we calculated the difference in the best decoding performance relative to PSID (**figure 6(b)**). For position decoding, ridge regression showed significantly lower *R^2^*values (mean difference = −0.09 ± 0.009 SEM, FDR-corrected Wilcoxon signed-rank test, *p* = 0.0012), Wiener filter (0.03 ± 0.004, *p* = 0.0012), MLP (0.04 ± 0.016, *p* = 0.042), CNN (0.20 ± 0.014, *p* = 0.0012), and GRU (0.24 ± 0.021, *p* = 0.0012) showed significantly higher performance. For velocity decoding, the other decoders exhibited a pattern similar to that observed for position decoding (all *p* = 0.001), except that the MLP did not show a significant increase in decoding performance. For comparison, we also evaluated the decoding performance of the SBP and 200–400 Hz power (**figure 6(c)**). For SBP, ridge regression (−0.15 ± 0.014), Wiener filtering (−0.01 ± 0.005), and MLP (−0.02 ± 0.018) showed lower position *R^2^* values than PSID, whereas CNN (0.15 ± 0.009) and GRU (0.22 ± 0.012) achieved higher performance. For 200–400 Hz power, all decoders except GRU (0.01 ± 0.025) showed lower *R^2^* than PSID. The velocity decoding exhibited a similar pattern. Overall, all three neural signals generally benefited from nonlinear decoders, with the LMP showing the greatest improvement.

Notably, different decoders achieved the best performance at different low-pass cutoff frequencies. Except for MLP, all decoders achieved their best position decoding performance with cutoff frequencies below 40 Hz. The median optimal cutoff frequencies were lowest for PSID and CNN, with a median of 5 Hz (95% CI: 5–8 and 5–10, respectively), followed by ridge regression (8 [5–40]), Wiener filter (8 [8–10]), and GRU (20 [20–30]). MLP performed best when using raw LMP without additional low-pass filtering (**figure 6(d)**). For velocity decoding, ridge regression,

PSID, CNN, and GRU all showed a median optimal cutoff frequency of 5 Hz, whereas the Wiener filter and MLP showed a slightly higher median of 8 Hz. Accordingly, most sessions exhibited higher *R^2^* values when using the optimal low-pass cutoff frequency than when using the raw LMP (**figure 6(e)**). The S-probe sessions exhibited a similar pattern, although the distribution of the optimal low-pass cutoff frequency was broader (**figure 7**). We further examined the optimal high-pass cutoff frequency for decoding and found that cutoff frequencies below 0.5 Hz were almost equally good for all decoders in both Utah-array and S-probe sessions (**figure 8**). Collectively, these findings suggest that both hypotheses are supported. The composite nature of LMP likely contributes to the superior performance of nonlinear decoders over linear decoders. However, the optimal low-pass cutoff frequency varied across decoders and even across recording sessions, highlighting the need for decoder- and session-specific optimizations.

**Figure 7.**
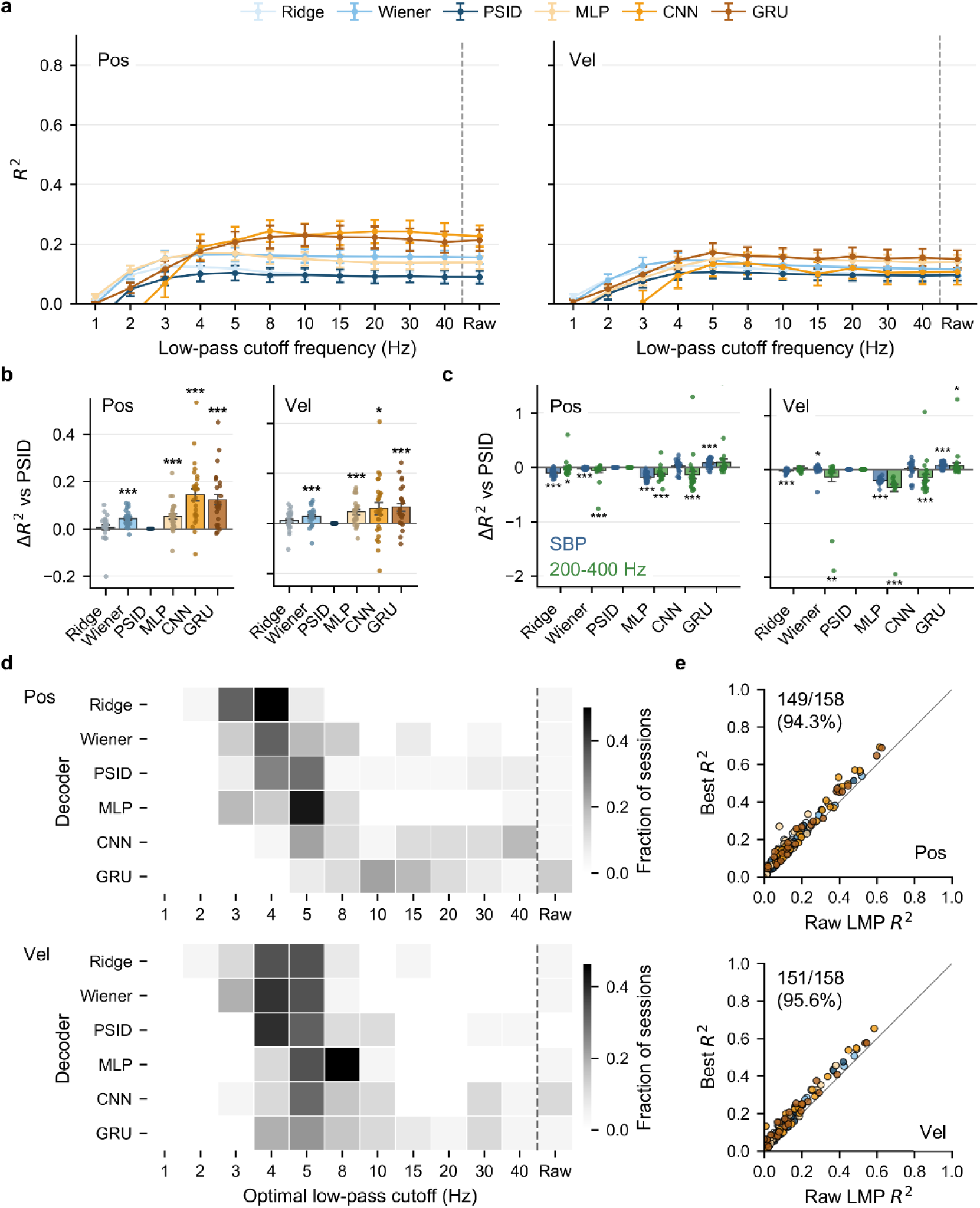
Evaluation of decoder performance and feature optimization in S-probe sessions. The legend is the same as in Figure 6. N = 26 recording sessions.

**Figure 8.**
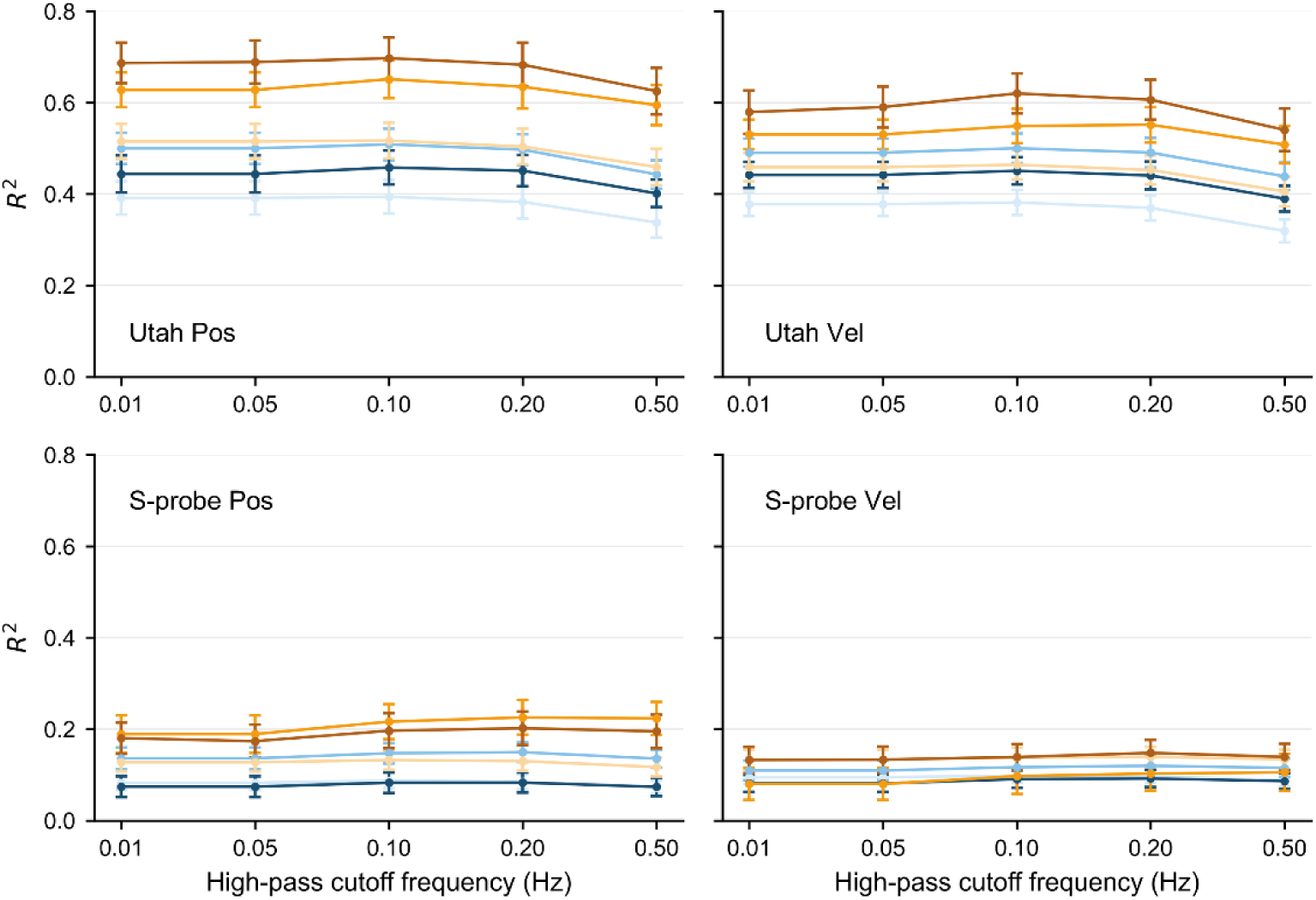
Effect of high-pass cutoff frequency on decoding performance. Decoding performance (*R*^2^) for position (Pos) and velocity (Vel) using Utah arrays (N = 11) and S-probes (N = 26) across different high-pass cutoff frequencies (0.01–0.50 Hz).

## 4. Discussion

Our study compared the motor decoding performance of different broadband signals recorded from the motor cortices of macaque monkeys using two types of electrode arrays: a planar Utah array and linear S-probes. We found that signals containing spiking activity or high-frequency components, such as SBP and high-gamma power, showed decoding performance that scaled with the number of recorded single units. In contrast, LMP, a low-frequency signal, achieved higher *R^2^*values in sessions recorded with Utah arrays than with S-probes, independent of single-unit count. Consistent with these decoding differences, the LMP exhibited lower inter-electrode correlation and greater variability in the preferred directions (PDs) in the Utah-array sessions. We further characterized the spectral and temporal properties of the LMP and found that its dominant decoding components overlapped substantially with the delta band (0.1–4 Hz) and movement-related cortical potentials (MRCPs). Notably, the optimal frequency range for LMP decoding (0.1– 8 Hz) was considerably lower than that of the raw LMP, suggesting that its decoding performance is primarily driven by its low-frequency components. Finally, LMP decoding benefited from nonlinear decoders, with CNN and GRU models outperforming the conventional linear approaches. Together, these findings suggest that LMP decoding depends more strongly on the spatial organization and temporal dynamics of low-frequency population activity, which may be better captured by dense planar sampling, highlighting the importance of electrode geometry and nonlinear temporal structure in neural decoding.

### 4.1. Horizontal versus vertical organization

The spatial heterogeneity of the LMP may arise from two distinct forms of organization within the motor cortical circuits along the horizontal and vertical axes. Horizontally, motor-related cortical areas are functionally specialized and spatially structured. Graziano et al. proposed the behavioral repertoire hypothesis, suggesting that the motor cortex represents a high-dimensional repertoire of behaviorally relevant actions that are mapped onto the two-dimensional cortical sheet while preserving local continuity (Graziano, Taylor and Moore, 2002; Graziano and Aflalo, 2007).

Consistent with this view, M1 exhibits a patchy, functionally specific intrinsic connectivity that closely follows its somatotopic organization (Card and Gharbawie, 2020). Moreover, Rubino, Robbins, and Hatsopoulos (2006) showed that beta-band oscillations propagated as traveling waves across M1 and PMd during motor preparation, with their propagation patterns carrying information about the upcoming movement (Rubino, Robbins and Hatsopoulos, 2006). Finally, a PMd–M1 beta gradient, characterized by greater high-beta activity in the PMd and stronger low-beta activity in the M1, further demonstrates that LFP properties vary systematically across cortical areas (Nougaret *et al*., 2024). Our results showed that interchannel correlations were generally lower in planar Utah arrays than in linear S-probes across broadband signals, except for SBP and high-gamma power, which contain spiking activity. Together, these findings suggest that the horizontal organization of the motor cortex forms a motor-related information-rich spatial architecture.

Vertically, neuronal populations are organized into laminae, supporting structured input– output information flow in the motor cortex. Superficial layers of M1 (L2/3) receive substantial somatosensory and premotor inputs and exhibit more choice-related activity, whereas deep layers (L5/6) give rise to extensive corticofugal and corticospinal projections containing more peri-movement and movement-related neurons (Weiler *et al*., 2008; Chandrasekaran *et al*., 2017; Persichetti *et al*., 2020). Therefore, cortical depth substantially influences the information captured by the electrodes. Although both superficial and deep layers encode movement and direction, neurons in deeper layers are more likely to encode directional information (Parikh, Marzullo and Kipke, 2009). Therefore, sampling of certain cortical layers may provide more effective information for neural decoding. Because Utah arrays primarily sample neurons around layer 5, whereas linear arrays span multiple cortical layers with only a subset of electrodes located in deeper layers, Utah arrays may provide a greater proportion of electrodes sampling information-rich neuronal populations despite using the same number of channels in the decoding analyses.

Given the fixed implantation depth of the Utah arrays, we could not determine whether other layers would provide comparable decoding performance to that observed for layer 5. Future studies using customizable or depth-adjustable planar arrays could address this question.

In addition, Mendoza-Halliday et al. (2024) further showed laminar differences in the LFP spectral composition, with higher-frequency activity predominating in superficial layers and low-frequency and alpha–beta activity becoming more prominent in deeper layers. Consistent with this, depth-dependent differences in intrinsic timescales are more pronounced in LFP than in SUA, whereas SUA exhibits clearer areal differences between M1 and PMd (Nandi *et al*., 2026). We found that the PDs were less diverse and more strongly correlated across the cortical laminae for low-frequency signals than for high-frequency signals recorded with linear arrays, suggesting distinct propagation properties across frequency components. Low-frequency signals, such as the LMP, were more homogeneous across cortical layers, potentially resulting in more redundant and less dynamically varying neural information recorded with a linear array. Overall, the variability in decoding performance across array configurations may reflect differences in both the horizontal sampling location and cortical depth.

### 4.2. Potential relationship between LMP and MRCP

Despite its strong behavioral relevance, the physiological origin of the LMP remains unclear. In our study, we found that the LMP exhibited fingerprint-like characteristics, with highly consistent direction tuning patterns across recording sessions within the same subject (**figure 4**). One potential explanation is the subject-specific organization of the motor cortical representations, could lead to stereotyped patterns of low-frequency population activity within an individual.

Consequently, the sequential activation of motor representations and thus the propagation of low-frequency waves may be highly stereotyped within each subject. In addition, arrays located in different brain areas may exhibit different PDs, which may be related to differences in implantation depth, as suggested by surgical observations. Whether LMP PDs vary across cortical layers remains an open question for future investigation.

The potential relationship between LMP and MRCP has received relatively limited attention. Two observations from previous studies are particularly relevant to this possibility. The first occurred in the study by Schalk et al. (2007), who introduced the LMP for the first time. In their figure 2, the channels showing strong correlations with movement parameters—the observation that motivated the introduction of the LMP—exhibited features highly reminiscent of typical MRCPs. However, because their analysis focused on frequency bands above 8 Hz, the potential correspondence between the 0.1–5 Hz component and LMP was not examined. The second occurred in a study by Gallego-Carracedo et al. (2022). In their figure 2F, the raw 0.5–4 Hz component and LMP were plotted together, and the two signals appear remarkably similar.

Nevertheless, their subsequent analyses used LFP power rather than the low-frequency signal itself, and they observed differences in decoding performance between the 0.5–4 Hz band power and LMP, which were easily interpreted as distinct neural components at that time. It should be noted that these retrospective analyses are not intended to suggest that previous studies overlooked this relationship, as those studies were not designed to investigate the origin of LMP. Rather, they provided additional retrospective evidence supporting the relationship between LMP and MRCP.

Our results showed that low-pass-filtered LMP components with optimized cutoff frequencies yielded better decoding performance than LMP, MRCP, or canonical frequency bands, such as the delta band (**figure 6**). These findings suggest that low-frequency neural activity is a highly dynamic signal, whose optimal frequency range may vary depending on multiple factors, including the recording area, noise, and behavioral context. Although we cannot yet determine what drives these dynamics or their physiological significance, they provide an interesting direction for future research. Overall, these findings highlight the importance of customizing the feature selection for motor decoding.

### 4.3. Implications for future BCI design

Given the rapid development of BCI technologies, particularly fully implanted wireless BCIs, electrode configuration has important implications for both the design of wireless communication modules and the selection of neural signals for online decoding (Guenther *et al*., 2009; Schwarz *et al*., 2014; Benabid *et al*., 2019; Rapeaux and Constandinou, 2021; Simeral *et al*., 2021; Mitchell *et al*., 2023; Patrick-Krueger, Burkhart and Contreras-Vidal, 2025). In the pursuit of higher channel density and reduced invasiveness, flexible linear arrays have increasingly been adopted as an alternative electrode configuration for intracortical BCI (Musk, 2019; Oh *et al*., 2025). This raises an important question for fully implanted systems: which neural signal components need to be transmitted for effective online decoding? Our results showed that decoding performance of low-frequency signals, such as the LMP, was strongly influenced by electrode configuration, whereas broadband signals containing spiking activity, such as SBP and high-gamma activity, were less dependent on electrode configuration and were more closely related to the availability of single-unit activity. These findings suggest that the optimal neural data to be transmitted may depend on the electrode configuration. For linear arrays, where a large number of recording channels can provide sufficient single-unit activity, transmitting TCRs or high-frequency signals may be sufficient for online decoding. In contrast, for planar arrays, both low-frequency signals, such as LMP, and high-frequency signals containing spiking activity may need to be retained to provide complementary information and compensate for potential loss of single-unit activity. Although the present study did not directly assess the long-term stability of LMP-based decoding, future longitudinal studies will be important to determine whether the observed configuration-dependent differences are maintained over extended recording periods.

Regarding optimal electrode configurations for decoding, a recent study suggested that motor cortical representations are spatially heterogeneous, yet functionally organized, with spatially distributed neurons exhibiting coordinated dynamics and shared task representations (Canfield *et al*., 2026). This organization highlights the need for flexible high-density probes capable of sampling neural activity across both cortical surface locations and depths. Several technologies have enabled such flexible three-dimensional sampling. Feingold et al. (2012) developed the Chronic Independently Movable Electrode (ChIME) system, which enabled chronic recording from up to 127 independently movable electrodes across multiple brain regions in awake macaques (Feingold *et al*., 2012). Voigts et al. (2013) developed flexDrive, allowing independent depth adjustment of up to 16 electrodes for chronic ensemble recordings in freely moving mice (Voigts *et al*., 2013). More recently, Kim et al. (2025) introduced a flexible, depth-adjustable 3D multishank array with approximately 0.1-mm depth resolution (Kim *et al*., 2025). Together, these approaches not only enable the sampling of distributed behavior-relevant neural dynamics across cortical depths for efficient decoding but also allow electrodes to be repositioned after implantation to mitigate gliosis-related signal degradation and optimize unit isolation.

The increasing demand for fully implantable and wireless BCIs has highlighted the neural data transmission bandwidth and power consumption as critical constraints for scalable neural recording. Conventional wireless intracortical BCIs often transmit broadband neural signals at high sampling rates, preserving the flexibility to extract spikes, LFPs, and other neural features offline but at the cost of high data rates and energy consumption, particularly as the channel count increases (Rizk *et al*., 2009). To address this limitation, alternative approaches perform on-implant signal processing and transmit more compact neural features such as TCRs, spike counts, and SBP, thereby substantially reducing wireless data and power consumption (Rizk *et al*., 2009;

Bullard *et al*., 2019; Even-Chen *et al*., 2020; Simeral *et al*., 2021). More recent architectures have further moved the decoding process onto the implant, transmitting only high-level outputs, such as position, velocity, or movement intention (Yan *et al*., 2024; Yin, Liu and Kornysheva, 2025).

Although this minimizes transmission demands, it limits access to underlying neural activity and flexibility for subsequent decoding or reanalysis. Against this background, our findings suggest an additional opportunity to reduce the transmission burden while retaining the neural signals for flexible decoding. Specifically, we found that the decoding performance could be optimized using substantially lower-frequency components than those typically transmitted or analyzed, whereas strong inter-electrode correlations, together with our channel-selection analysis, suggest that only a subset of channels may be sufficient to achieve optimal decoding performance. Thus, frequency- and channel-selective neural feature transmission could further reduce wireless data rates and power consumption without requiring the decoder to be fully implemented on the implant.

## Data availability statement

The dataset recorded using linear arrays is publicly available through Zenodo (DOI: https://doi.org/10.5281/zenodo.14196227). The dataset recorded using Utah arrays is available from the authors upon reasonable request.

## Acknowledgements

The authors would like to thank Yingming Wei, Yiheng Zhang, and Ruichen Zheng for their assistance with the surgeries, and Qifan Wang for animal training and guidance in neural data collection. This work was supported by grants from the Brain Science and Brain-like Intelligence Technology—National Science and Technology Major Project (2026ZD0220300) and Major Program of National Natural Science Foundation of China (T2594604033) to H.C and Shanghai Postdoc Fellowship Funding (Y95CN51621) to C.Z.

## Code availability

Code for reproducing figures is available at https://github.com/zhengcong26/Electrode2types_motor_decoding.

## Author contributions

C.Z. and H.C. conceived of and designed the experiments. C.Z. and Y.X. performed experiments.

C.Z. analyzed the data and wrote the manuscript with input from all authors. H.C. and C.Z. acquired the funding. H.C. supervised the project.

